# *Botrytis cinerea* infection reshapes the grape berry microbiome during noble rot

**DOI:** 10.64898/2026.05.29.728822

**Authors:** Dario Cantu, Barbara Blanco-Ulate, Greg Allen, Rosa Figueroa-Balderas

**Author notes:** Corresponding author: D. Cantu.

## Abstract

Noble rot, caused by *Botrytis cinerea*, profoundly alters grape berry physiology and is essential to produce botrytized wines. In this study, we profiled bacterial and fungal communities associated with *Vitis vinifera* cv. Sémillon berries across four stages of noble rot development and four consecutive vintages using 16S rRNA gene and ITS1 amplicon sequencing. Noble rot stage significantly impacted the structure of bacterial communities across vintages, while fungal communities showed more variable vintage-dependent responses. Bacterial alpha diversity increased consistently at advanced stages of infection (S3), coinciding with a marked shift from *Pseudomonas*-dominated communities toward acetic acid bacteria, particularly *Gluconobacter*, which was significantly enriched at S3 in all vintages. Fungal communities remained dominated by Sclerotiniaceae throughout infection, consistent with pervasive *B. cinerea* colonization, while non-*Botrytis* fungal taxa shifted from filamentous fungi such as *Cladosporium* and *Alternaria* toward fermentative yeasts including *Hanseniaspora* and *Lachancea*. Co-occurrence network analysis revealed a positive association between *Gluconobacter* and these fermentative yeasts, suggesting coordinated enrichment of oxidative and fermentative microorganisms at advanced noble rot stages. Together, these results reveal a reproducible stage-associated microbial succession during noble rot progression and identify acetic acid bacteria as consistent markers of advanced infection.

## Introduction

*Botrytis cinerea* is among the most economically damaging fungal pathogens in viticulture worldwide, causing gray bunch rot and substantial yield and quality losses across wine-growing regions. Under specific microclimatic conditions, however, characterized by alternating humid nights and dry sunny days, the same pathogen produces noble rot, an infection process that promotes favorable metabolic and compositional changes in ripe berries, which are essential for the production of botrytized dessert wines, including Sauternes, Tokaj Aszú, and Trockenbeerenauslese (Blanco-Ulate et al. 2015; Magyar and Soós 2016).

Noble rot causes profound physicochemical alterations in the grape berry, including skin cracking, dehydration, increase in sugar concentration, shifts in organic acid composition (Blanco-Ulate et al. 2015; Magyar 2011; Ribéreau-Gayon et al. 2006), and accumulation of gluconic acid from both fungal and bacterial activity (Barata et al. 2012; Magyar and Soós 2016). We previously showed that noble rot triggers extensive metabolic reprogramming in Sémillon berries, activating stress responses and ripening-associated pathways that are otherwise only weakly expressed during normal ripening of white-skinned berries (Blanco-Ulate et al. 2015). These physiological and chemical changes occur within a microbially rich environment. Late harvest ripe grape berries are colonized by diverse communities of fungi, yeasts, and bacteria (Barata et al. 2012), and the physiological and metabolic alterations associated with *B. cinerea* infection can substantially reshape the berry-associated microbial niche. (Barata et al. 2012; Bokulich et al. 2014)

The grape berry microbiome has received increasing attention over the past decade, driven in part by its recognized role in shaping fermentation dynamics, aroma compound production, the microbial basis of regional wine character, and spoilage risk (Barata et al. 2012; Bokulich et al. 2014; Griggs et al. 2021). Amplicon sequencing studies targeting fungal ITS regions and bacterial 16S rRNA genes have shown that ripening berry surfaces harbor structured microbial communities whose composition shifts progressively through the season, with a marked transition at veraison coinciding with sugar accumulation and cuticle softening (Kecskeméti et al. 2016; Liu and Howell 2021). As ripening advances, these physiological changes favor the proliferation of fermentative yeasts, particularly *Hanseniaspora* and *Metschnikowia* spp., which become core members of the berry microbiota at harvest and seed the early stages of wine fermentation (Barata et al. 2012). The composition of the microbial communities is shaped by regional, varietal, climatic, and management factors, with geographic origin emerging as a major determinant of microbiome structure, a phenomenon described as microbial terroir (Bokulich et al. 2014; Gilbert et al. 2014; Griggs et al. 2021).

More recently, the host genome itself has been implicated in microbiome assembly, with loci associated with plant immune responses influencing the abundance of pathogenic fungi, including *Botrytis* spp., as well as fermentative yeasts such as *Saccharomyces cerevisiae* (Flörl et al. 2025), suggesting that the genetic determinants of disease susceptibility and microbiome composition are partially intertwined. Acetic acid bacteria (AAB), primarily *Acetobacter* and *Gluconobacter* spp., occur at low abundance on healthy grape berries but proliferate extensively on damaged fruit, including *Botrytis*-affected berries, where populations may reach ~10^6^ CFU/g or higher (Barata et al. 2012). Studies of grape sour rot have consistently identified acetic acid bacteria, including *Acetobacter* spp., as enriched in diseased tissue relative to asymptomatic berries (Hall et al. 2018). More broadly, pathogen infection reshapes phyllosphere microbial communities across plant species. Rust fungus infection in *Malus* reduced bacterial alpha diversity while increasing fungal alpha diversity during lesion expansion (Zhang et al. 2023), whereas *Diaporthe citri* infection in citrus reduced community evenness, introduced new taxa, and increased network complexity (Li et al. 2022).

Upon *Botrytis* infection, berries become co-colonized by bacteria and yeasts that begin modifying the berry chemistry before harvest (Magyar 2011). However, how these microbial communities change during noble rot progression remains largely uncharacterized. Bacterial diversity in botrytized wine fermentations was reported to be substantially higher than in fermentations of healthy grapes harvested from the same vineyard, and AAB were strongly suppressed following *Saccharomyces* inoculation during fermentation (Bokulich et al. 2012), directly linking pre-harvest microbiome assembly to microbial activity during vinification and wine composition.

Here, we characterize the bacterial and fungal microbiomes of naturally-infected *Vitis vinifera* cv. Sémillon berries collected across four vintages and four stages of noble rot progression, from asymptomatic berries to heavily colonized sporulating fruit. Using 16S rRNA gene and ITS1 amplicon sequencing, we show that *B. cinerea* infection drives a reproducible, stage-structured microbial succession marked by increasing bacterial diversity, enrichment of AAB, particularly *Gluconobacter*, within the S3 core microbiome, and a shift in the non-*Botrytis* fungal community from filamentous fungi toward fermentative yeasts. Co-occurrence network analysis further reveals a positive association between *Gluconobacter* and fermentative yeasts, including *Hanseniaspora* and *Lachancea*, at advanced infection stages, pointing to the coordinated assembly of an oxidative-fermentative microbial community with direct implications for botrytized wine production.

## Material and Methods

### Biological material and sampling

Ripe *Vitis vinifera* cv. Sémillon berries were harvested from the Dolce Winery vineyards in Napa, California across four consecutive vintages: November 6, 2012, October 13, 2013, November 6, 2014, and November 20, 2015. Harvest dates were chosen to coincide with the days selected by the Dolce winemaker for botrytized wine production. All sample collections were performed between 8 and 10 AM. Weather conditions at the time of collection were: 17.2 ± 3.4°C with 68.3 ± 12.1% relative humidity (2012), 10.6 ± 5.1°C with 83.0 ± 14.7% relative humidity (2013), 14.4 ± 3.1°C with 78.0 ± 15.0% relative humidity (2014), and 8.2 ± 3.4°C with 93.8 ± 8.5% relative humidity (2015).

Three stages of noble rot progression (S1, S2, S3) were defined based on visual assessment of symptom severity and were confirmed by measuring *B. cinerea* biomass (Blanco-Ulate et al., 2015). Briefly, S1 berries showed an initial color change from yellow to pink; S2 berries had softened skin with dark pink coloration; S3 berries were fully colonized with evident mycelial growth and sporulation. Ripe berries without visible *B. cinerea* infection were sampled simultaneously as asymptomatic controls. Samples were collected randomly from vines in the same rows across all vintages to minimize spatial confounding effects. Individual berries were harvested from clusters at different positions within the vine canopy to account for variation in sun exposure. For microbiome profiling, three biological replicates per stage per vintage were collected, each consisting of an independent pool of 20 berries from at least five different vines. Samples were transported on ice and stored at −80°C.

### DNA extraction and library preparation

Genomic DNA was extracted at room temperature using a modified CTAB-based protocol. Briefly, berries samples were ground to a fine powder in liquid nitrogen and incubated in pre-warmed extraction buffer (65°C) supplemented with 2% (v/v) 2-mercaptoethanol and Proteinase K (0.4 mg/ml). Following incubation at 65°C for 30 minutes, samples were centrifuged, and the supernatant was extracted twice with chloroform:isoamyl alcohol (CIA, 24:1, v/v), including an RNAse A (0.2 mg/ml) treatment between extractions. Genomic DNA was precipitated using sodium acetate and isopropanol, pelleted by centrifugation, and resuspended in Qiagen EB buffer (Qiagen, USA). Genomic DNA was further purified using sequential phenol:CIA (25:24:1) and CIA extractions, followed by selective polysaccharide removal using low-volume ethanol precipitation (0.3X volume). Genomic DNA was then precipitated with 1.7X volume of ethanol, washed with 70% ethanol, air dried, and resuspended in warmed water. DNA quality was assessed by spectrophotometry and agarose gel electrophoresis, and concentration was quantified using a Qubit fluorometer (Thermo Scientific, USA).

DNA was amplified using a unique 8 nt barcode primer set for 16S and ITS. For the 16S rRNA gene V4 region we used primers 515F/806R (Walters et al. 2015) and for the fungal ITS1 region we used BITS/B58S3 primers (Bokulich and Mills 2013). PCR was developed in a 25 µl PCR reaction using GoTaq polymerase (Promega Corporation, USA) in a Verity thermal cycler (Applied Biosystems, USA) under the following conditions: an initial denaturation at 95°C for 45 seconds, followed by 35 cycles at 95°C for 1 minute, 55°C for 1 minute, and 72°C for 1 minute, with afinal extension at 72°C for 10 minutes and a hold at 4°C after. Amplicon size was verified by electrophoresis, and PCR products were purified using Ampure XP magnetic beads (Roche, USA). DNA concentration for each purified amplicon was determined using a Qubit fluorometer (Thermo Scientific, USA). Equimolar amounts of all barcoded amplicons were pooled into a single library. A total of 500 ng pooled DNA was end-repaired, A-tailed and ligated to single index adapter (Kapa Biosystems, USA). Following adapter ligation, two consecutive 1X bead-based cleanups were performed, and library concentration and size distribution were assessed using a Qubit fluorometer and Bioanalyzer (Agilent Technologies, USA), respectively. Amplicon libraries were sequenced in 250 bp paired-end mode on an Illumina MiSeq platform at the DNA technologies Core, University of California, Davis.

### Bioinformatic analysis of amplicons

Raw paired-end reads were processed using FASTX Toolkit (v0.0.14; Gordon and Hannon 2010). Barcodes were extracted from the first 8 bases of R1 reads using fastx_trimmer, and reads were trimmed to remove the first 8 bases prior to demultiplexing. Samples were demultiplexed using QIIME 2 (v2026.4; Bolyen et al. 2019) with the qiime demux emp-paired command with Golay error correction disabled, using 8-nucleotide barcodes. The two 16S sequencing runs shared the same samples and were processed independently; ASV tables and representative sequences were merged by summing read counts. The same procedure was applied to the two ITS runs.

Denoising and chimera removal were performed with DADA2 (Callahan et al. 2016) as implemented in QIIME 2, using paired-end mode with no truncation. Bacterial ASVs were taxonomically classified using a Naive Bayes classifier trained on the SILVA 138 99% OTU reference database trimmed to the 515F/806R V4 region. Fungal ASVs were classified against the UNITE v10.0 (February 2025) 99% reference database. Sequences assigned to chloroplasts or mitochondria were removed from the 16S dataset. A phylogenetic tree for bacterial ASVs was constructed using the qiime phylogeny align-to-tree-mafft-fasttree pipeline.

### Alpha diversity

Alpha diversity was calculated on the full ASV tables. Analyses were performed on non-rarefied ASV tables to avoid discarding valid sequencing reads (McMurdie and Holmes 2014). For bacteria, Shannon entropy, observed ASVs, Chao1 richness, Faith’s phylogenetic diversity, and Pielou’s evenness were calculated using QIIME 2. For fungi, the same metrics were calculated except for Faith’s phylogenetic diversity, which requires a phylogenetic tree. Diversity metrics were calculated separately for each vintage year (2012, 2013, 2014, 2015). Due to insufficient sequencing depth (fewer than 5,000 reads), the 2015 S3 bacterial samples were excluded from downstream analyses. Differences in alpha diversity across noble rot stages were tested using the Kruskal-Wallis test, and pairwise comparisons were performed using the Wilcoxon rank-sum test (p<0.05) in R (v4.5).

### Beta diversity

Beta diversity was calculated on the full ASV tables without rarefaction. For bacteria, weighted UniFrac and unweighted UniFrac distances were calculated using the qiime diversity beta-phylogenetic command in QIIME 2, using the rooted phylogenetic tree constructed from bacterial ASVs. For fungi, Bray-Curtis dissimilarity was calculated using qiime diversity beta. All metrics were calculated separately for each vintage year to avoid confounding effects of year on community composition. Principal coordinate analysis (PCoA) was performed on each distance matrix using qiime diversity pcoa. Differences in community composition across noble rot stages were tested using PERMANOVA with 999 permutations implemented in qiime diversity beta-group-significance.

### Taxonomic composition

Bacterial and fungal ASV tables were collapsed to genus level (taxonomic level 6) using the qiime taxa collapse command in QIIME 2. Relative abundances were calculated per sample and the top 25 genera by mean relative abundance across all samples were retained; remaining genera were grouped as “Other”. For fungal communities, *Sclerotiniaceae* were excluded prior to genus-level visualization to allow inspection of the remaining community, given the overwhelming dominance of this family across all samples and stages. A separate family-level visualization including Sclerotiniaceae was produced using ASV tables collapsed to taxonomic level 5.

### Differential abundance analysis

Differential abundance of bacterial and fungal genera across noble rot stages was tested using ANCOM-BC (Analysis of Compositions of Microbiomes with Bias Correction) as implemented in the qiime composition ancombc plugin in QIIME 2. Analyses were performed separately for each vintage year using genus-level collapsed feature tables, with control samples as the reference level and noble rot stage as the model formula. Taxa with a false discovery rate-corrected q-value < 0.05 were considered significantly differentially abundant. Differential abundance analysis for fungal communities could not be performed for 2012 and 2013 due to the absence of fungal reads in control samples in 2012 and insufficient control sample size in 2013, consistent with the low fungal biomass prior to *Botrytis cinerea* infection in those vintages.

### Acetic acid bacteria profiling

Acetic acid bacteria were identified by filtering the bacterial genus-level feature table to retain only taxa assigned to the family Acetobacteraceae, including *Gluconobacter, Komagataibacter, Acetobacter, Kozakia, Gluconacetobacter*, and *Asaia*. Relative abundances were calculated per sample as a proportion of total bacterial reads. Total AAB abundance per sample was computed by summing the relative abundances of all AAB genera. Mean relative abundance per genus per stage and year was calculated across biological replicates.

### Co-occurrence network analysis

Co-occurrence networks were constructed using Spearman rank correlations calculated on genus-level relative abundance matrices for the 58 samples with both 16S and ITS data. Genera retained for network analysis were present in at least 30% of samples with a mean relative abundance of at least 0.01%. Correlation coefficients and associated p-values were calculated using the corr.test function from the psych R package (v2.6.3), with Benjamini-Hochberg correction for multiple testing. Bacterial within-kingdom correlations were filtered at |r| ≥ 0.5 and q < 0.05. Cross-kingdom correlations between bacterial and fungal genera were filtered at |r| ≥ 0.3 and q < 0.05. Networks were constructed and visualized using the igraph (v2.3.1) and ggraph (v2.2.2) R packages, with the Fruchterman-Reingold layout algorithm. Node size was scaled to mean relative abundance and edge width to correlation strength.

## Results

### Sampling and data collection

Grape berries of *V. vinifera* cv. Sémillon displaying noble rot symptoms were sampled across four vintages (2012, 2013, 2014, and 2015) from the same commercial vineyard, following the criteria of Blanco-Ulate et al. (2015). Briefly, *B. cinerea* infections were categorized into three stages of noble rot development (**Fig. 1**). The initial stage of infection (S1) was characterized by a color change of the berries from yellow to pink. As the infection progressed, the berry skin became softer and dark pink (S2). Berries at S3 (*pourri plein*) were fully rotten but not dry, with cracked skins, evident mycelial growth and sporulation, and purple-brown coloration. Berries showing no visible symptoms of infection were harvested simultaneously as asymptomatic controls (C).

**Figure 1.**
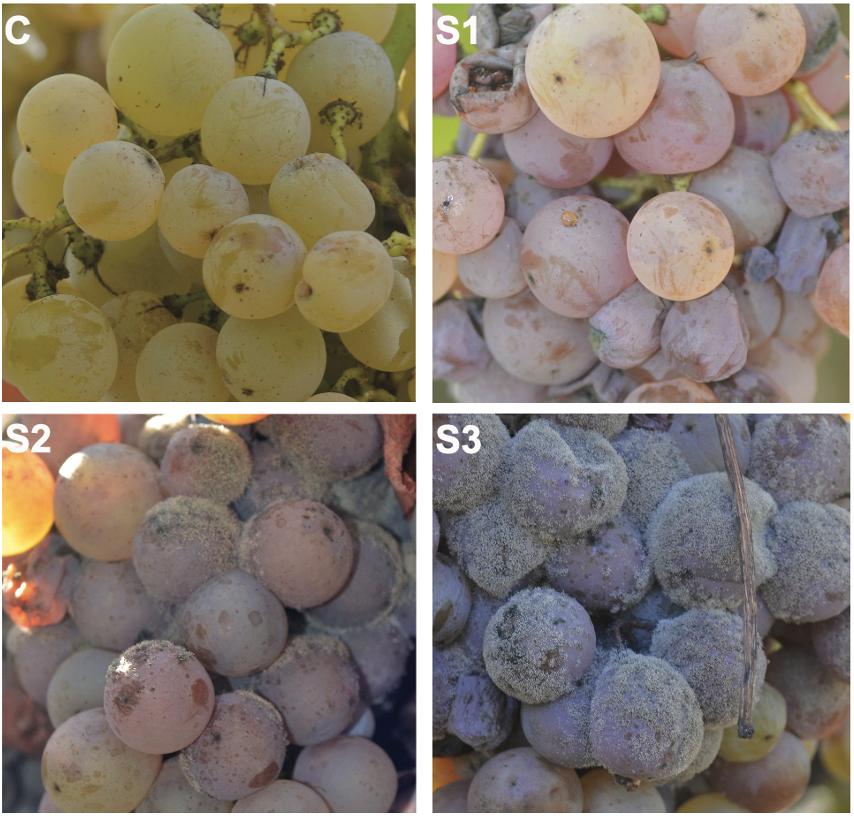
Stages of noble rot in *Vitis vinifera* cv. Sémillon. Representative grape clusters at the control stage (C, healthy berries with no visible *Botrytis cinerea* infection), early infection (S1, color change from yellow to pink with incipient mycelial growth), mid infection (S2, softened skin turning dark pink with visible sporulation), and advanced infection (S3, *pourri plein*, berries fully covered by dense fungal sporulation with extensive dehydration and purple-brown coloration). Staging following Blanco-Ulate et al. (2015).

Bacterial and fungal communities were profiled by sequencing the 16S rRNA gene V4 region and the ITS1 region, respectively, across 66 bacterial and 70 fungal samples representing four vintages and four noble rot stages. After quality filtering, denoising, and chimera removal, ten of twelve 2015 S3 bacterial samples were excluded due to insufficient sequencing depth (fewer than 5,000 reads per sample after denoising; mean of 2,393 reads, range: 658 to 4,633), leaving 66 bacterial samples for downstream analysis (**Supplementary Table S1**). A total of 1,269,688 high-quality bacterial reads were retained, yielding 2,088 unique bacterial ASVs, with a mean of 19,238 reads per sample (range: 4,342 to 40,374). For fungi, 6,098,321 reads were retained across 70 samples, yielding 3,344 unique fungal ASVs, with a mean of 87,119 reads per sample (range: 0 to 360,117).

### Bacterial alpha diversity increases with noble rot progression

Alpha diversity was assessed using Shannon entropy, observed ASVs, Chao1 richness, Faith’s phylogenetic diversity (bacteria only), and Pielou’s evenness; full results for all metrics are reported in **Supplementary Tables S2** (bacteria) and **S3** (fungi). Shannon entropy differed significantly across noble rot stages in bacterial communities in 2012 (Kruskal-Wallis, p = 0.042), 2013 (p = 0.050), and 2015 (p < 0.001), but not in 2014 (p = 0.123; **Fig. 2**). In 2012 and 2014, S3 showed the highest Shannon entropy (4.693 ± 0.297 and 5.641 ± 0.162, respectively), substantially higher than control, S1, and S2. In 2013, only control and S3 were sampled, and S3 showed markedly higher diversity (4.734 ± 0.569 vs. 3.783 ± 0.044 in control). In 2015, a stepwise increase was observed from control to S2, with S3 excluded due to insufficient sequencing depth. Pielou’s evenness mirrored the Shannon pattern, with significant stage effects in 2012 (p = 0.025) and 2013 (p = 0.050), driven by higher evenness at S3, while observed ASVs, Chao1 richness, and Faith’s phylogenetic diversity were significant only in 2015 (p < 0.001 for all three), where the larger sample size provided sufficient statistical power.

**Figure 2.**
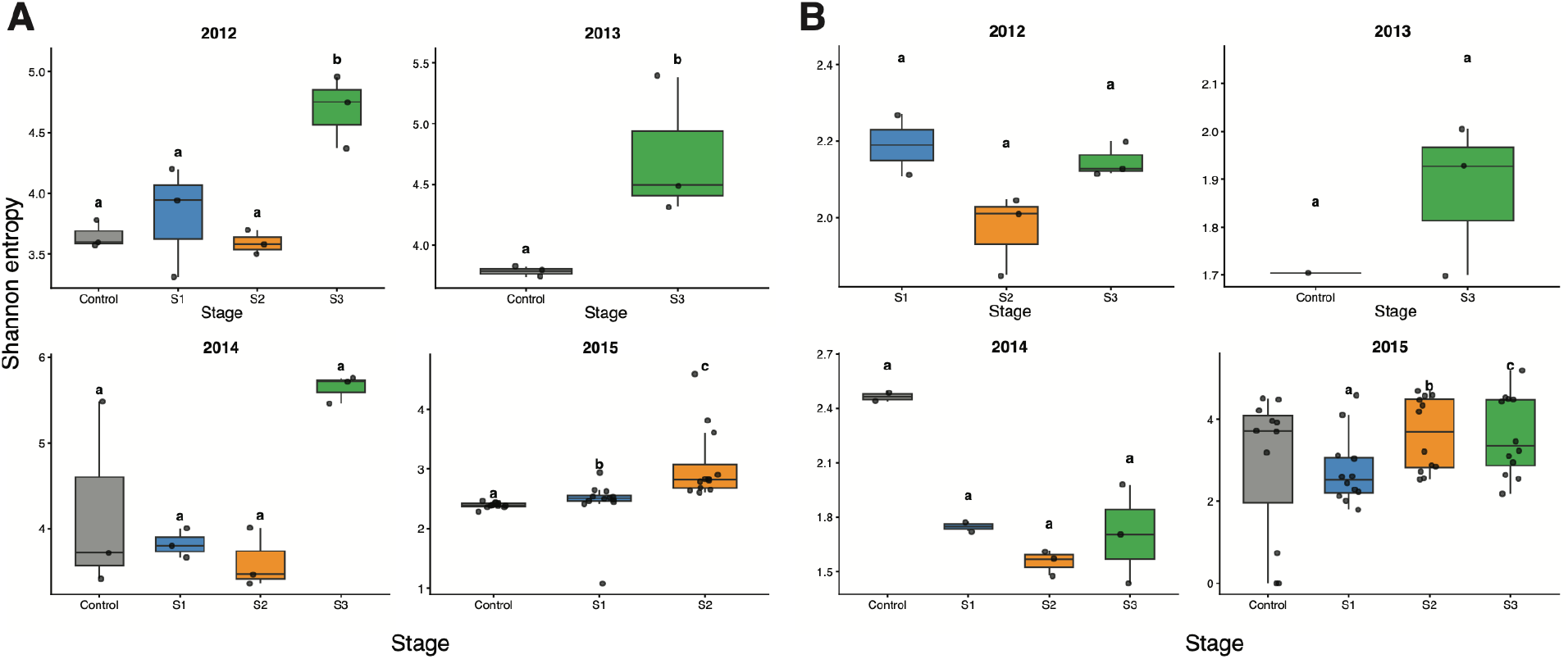
Alpha diversity (Shannon entropy) of bacterial (A) and fungal (B) communities across noble rot progression stages by year. Boxplots show the distribution of Shannon entropy at four noble rot stages: control (pre-infection), S1 (early infection), S2 (mid infection), and S3 (late infection) in 2012, 2013, 2014, and 2015. Individual data points are shown. Letters indicate statistically significant differences between stages based on pairwise Wilcoxon rank-sum tests (p<0.05). Note that 2013 and 2014 did not include all four stages. 2015 S3 bacterial samples with fewer than 5,000 reads were excluded due to insufficient sequencing depth.

For fungal communities, Shannon entropy did not differ significantly across noble rot stages in any year (**Supplementary Table S3**), although 2015 showed a trend toward higher diversity at S3 (p = 0.093). Fungal alpha diversity patterns were more variable across vintages and stages than those observed for bacteria, consistent with the high dispersion of fungal community composition in beta diversity analyses. Together, these results indicate that advanced noble rot stages are associated with increased bacterial alpha diversity, whereas fungal diversity responses are less consistent across vintages.

### Noble rot stage is associated with shifts in bacterial and fungal community composition

Principal coordinate analysis (PCoA) of weighted UniFrac distances revealed consistent separation of bacterial communities across noble rot stages (**Fig. 3A**). In all years with complete stage sampling (2012 and 2014), PC1 explained the majority of variance (70.4% and 78.7%, respectively), with S3 samples consistently separated from control, S1, and S2 along this axis. In 2013, where only control and S3 samples were available, the same separation was evident (PC1 = 77.7%). In 2015, where S3 samples were excluded due to insufficient sequencing depth, S1 and S2 samples clustered tightly while several S2 outliers were visible, suggesting greater community variability at mid-infection stages in that vintage.

**Figure 3.**
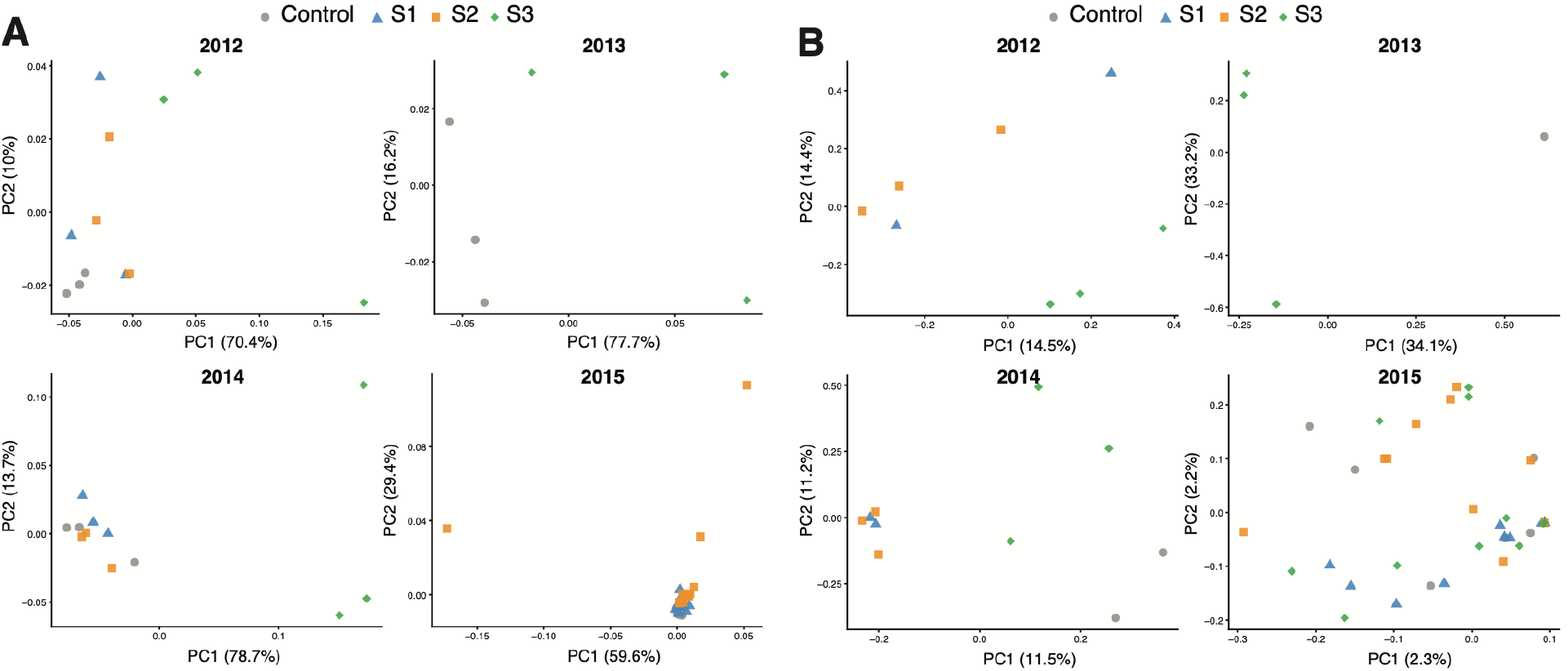
Beta diversity of bacterial and fungal communities across noble rot stages by year. Parincipal coordinate analysis (PCoA) of (A) weighted UniFrac distances for bacterial communities (16S rRNA gene V4 region) and (B) Bray-Curtis dissimilarity for fungal communities (ITS1 region), calculated separately for each vintage year (2012, 2013, 2014, 2015). Each point represents one sample, colored and shaped by noble rot stage: control (gray circles), S1 (blue triangles), S2 (orange squares), and S3 (green diamonds). The percentage of variance explained by each principal coordinate is shown on the respective axis. PERMANOVA results are reported in **Supplementary Table S4**. Note that 2013 includes only control and S3 samples; 2015 bacterial samples lack S3 due to exclusion for insufficient sequencing depth, and 2013 ITS includes only control and S3 samples.

Fungal community composition based on Bray-Curtis dissimilarity showed a different pattern (**Fig. 3B**). PC1 explained considerably less variance than in the bacterial analysis (14.5%, 34.1%, 11.5%, and 2.3% in 2012, 2013, 2014, and 2015, respectively), indicating greater overall variability in fungal communities. In 2012 and 2013, S3 samples tended to separate from earlier stages, while in 2014 S3 samples were more dispersed. In 2015, with all four stages represented and the largest sample size, communities showed substantial overlap among stages with no clear directional separation, suggesting higher inter-sample variability in fungal community composition within stages compared to bacteria.

To test whether noble rot stage significantly explained variation in microbial community composition, PERMANOVA with 999 permutations was applied to weighted and unweighted UniFrac distance matrices for bacteria and Bray-Curtis dissimilarity for fungi, calculated separately for each vintage year (**Supplementary Table S4**). PERMANOVA confirmed significant effects of noble rot stage on bacterial community composition in 2012 (weighted UniFrac: pseudo-F = 2.997, p = 0.009; unweighted UniFrac: pseudo-F = 1.586, p = 0.001), 2014 (weighted UniFrac: pseudo-F = 10.069, p = 0.015; unweighted UniFrac: pseudo-F = 1.529, p = 0.001), and 2015 (weighted UniFrac: pseudo-F = 1.403, p = 0.034; unweighted UniFrac: pseudo-F = 2.862, p = 0.001). In 2013, where only control and S3 samples were available, the stage effect was marginally non-significant for both metrics (weighted UniFrac: pseudo-F = 5.050, p = 0.107; unweighted UniFrac: pseudo-F = 1.565, p = 0.099), likely reflecting the low statistical power with only six samples.

For fungal communities, PERMANOVA revealed significant stage effects in 2012 (Bray-Curtis: pseudo-F = 1.010, p = 0.033) and 2014 (pseudo-F = 1.017, p = 0.003), but not in 2013 (pseudo-F = 1.034, p = 0.256) or 2015 (pseudo-F = 1.001, p = 0.320). The lack of significance in 2015 despite the largest sample size suggests high variability in fungal community composition within stages in that vintage, consistent with the dispersed PCoA ordination observed in **Fig. 3B**.

### *Pseudomonas* dominates early noble rot stages while acetic acid bacteria emerge at advanced infection

The bacterial community was dominated by *Pseudomonas* across all years and stages, accounting for the majority of reads in control, S1, and S2 samples (**Fig. 4A**; **Supplementary Table S5**). Community composition shifted markedly at S3, where *Pseudomonas* relative abundance decreased and a more diverse community emerged, with notable increases in *Gluconobacter, Komagataibacter*, and *Acetobacter. Acinetobacter* and unclassified taxa were consistently present as minor components across all stages. In 2014, S3 samples showed the most dramatic community shift, with *Gluconobacter* and *Komagataibacter* reaching their highest relative abundances. In 2015 samples, where S3 was excluded, the community was remarkably uniform across control and S1, with greater variability emerging at S2.

**Figure 4.**
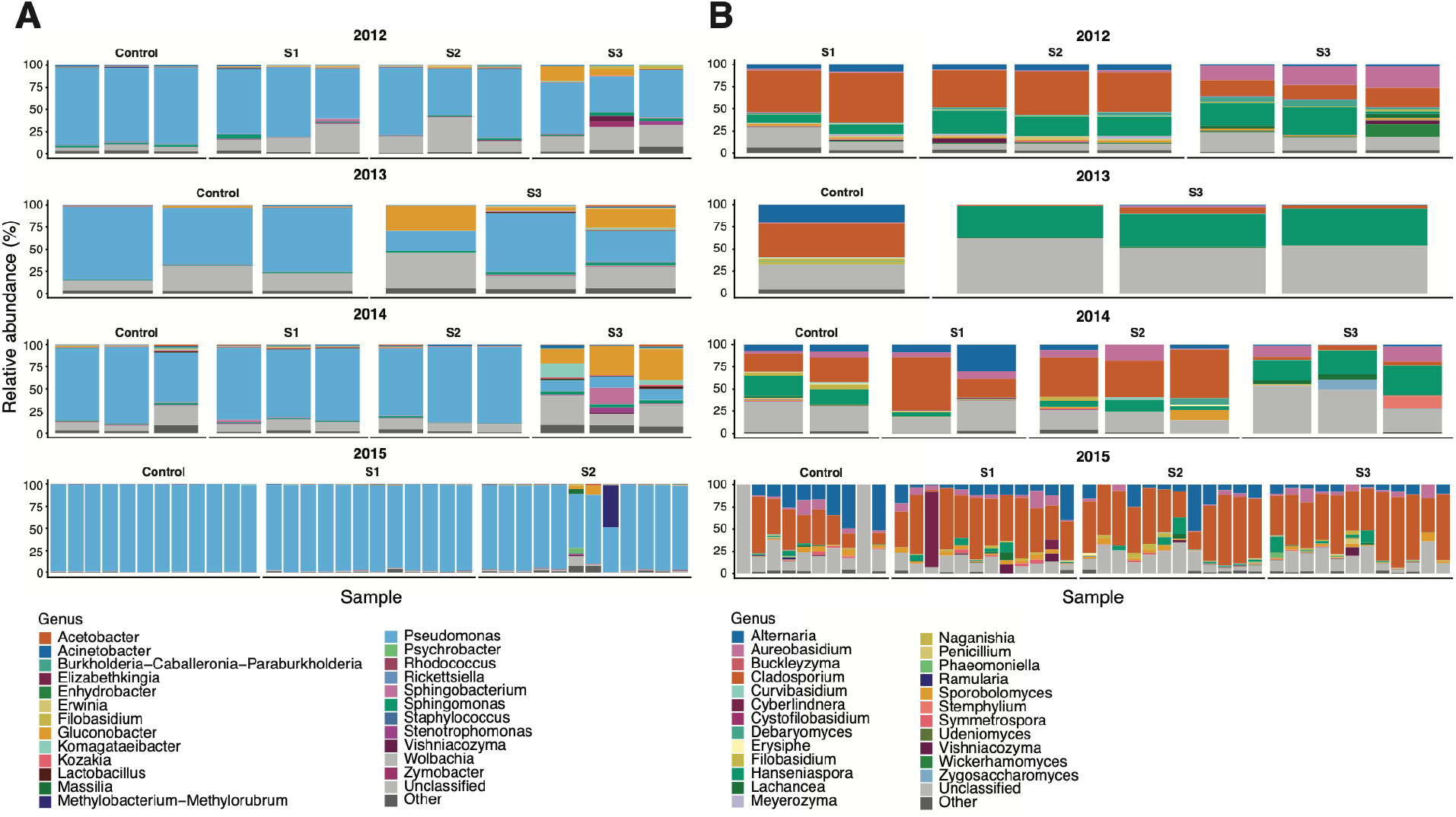
Relative abundance of bacterial (A) and fungal (B) communities at genus level across noble rot stages and vintages. Stacked barplots show the relative abundance (%) of the top 25 genera for bacteria (16S rRNA gene V4 region) and fungi (ITS1 region, Sclerotiniaceae excluded) in each sample, grouped by noble rot stage (Control, S1, S2, S3) and year (2012, 2013, 2014, 2015). Each bar represents one sample. Genera not in the top 25 are grouped as “Other”. Note that 2013 bacterial and fungal samples include only Control and S3 stages, 2015 bacterial samples lack S3 due to exclusion for insufficient sequencing depth, and fungal Sclerotiniaceae are excluded from panel B and shown separately in **Supplementary Figure 1**. Bacterial taxonomy was assigned using the SILVA 138 database; fungal taxonomy was assigned using the UNITE v10.0 database.

At the family level, Sclerotiniaceae dominated the fungal community across all years and stages, confirming the pervasive presence of *Botrytis cinerea* throughout noble rot progression (**Supplementary Fig. 1; Supplementary Table S6**). Sclerotiniaceae relative abundance was highest in 2012 and 2013, where it accounted for over 90% of reads in most samples. In 2015, Sclerotiniaceae remained the dominant family but with greater variability, and Cladosporiaceae showed higher relative abundance particularly in control samples, suggesting that *Cladosporium* is an early colonizer progressively displaced by *Botrytis cinerea* as infection advances. After excluding Sclerotiniaceae, the remaining fungal community was dominated by *Cladosporium* and *Alternaria* in early stages, with *Hanseniaspora* increasing at later stages (**Fig. 4B; Supplementary Table S7**). In 2012, *Cladosporium* dominated S1 and S2 while S3 showed increased diversity with *Aureobasidium, Hanseniaspora*, and multiple yeast genera. In 2013, the control was dominated by *Alternaria* while S3 showed increased *Cladosporium* and unclassified taxa. In 2014, *Cladosporium* dominated across all stages with *Hanseniaspora* emerging at S3. In 2015, the community was highly variable with *Cladosporium, Alternaria, Aureobasidium*, and *Hanseniaspora* all present across stages.

### *Gluconobacter* is consistently enriched at advanced noble rot stages while fungal communities shift from filamentous fungi toward yeasts

ANCOM-BC identified 47 unique bacterial genera significantly enriched or depleted relative to control across all years and stages (95 significant genus-stage-year combinations, q < 0.05; **Fig. 5A; Supplementary Table S8**). The most consistent finding across years was the enrichment of *Gluconobacter* at S3, which was significant in 2012, 2013, and 2014 (q < 0.05), with log fold changes increasing progressively from S1 to S3. *Erwinia* showed strong enrichment at S3 in 2012 and depletion in 2014, suggesting vintage-specific dynamics. *Elizabethkingia* was significantly enriched at S1 in 2012 but depleted at S3 in 2015, indicating an early colonizer that declines with infection progression. *Wolbachia, Hanseniaspora, Vishniacozyma*, and *Zymobacter* were enriched at S3 in 2012. In 2013, where only control and S3 were available, *Gluconobacter* and *Wolbachia* were significantly enriched at S3. In 2014, *Lactobacillus, Oenococcus*, and *Hymenobacter* were significantly depleted at S2 and S3, while *Gluconobacter* showed strong enrichment. In 2015, *Gluconobacter* remained the most significantly enriched taxon at S3, along with *Vishniacozyma* enrichment and *Pseudomonas* depletion. Notably, *Acetobacter* and *Komagataibacter*, the other main acetic acid bacteria genera, showed positive log fold changes at S3 across multiple years but did not reach statistical significance after FDR correction, likely reflecting high variability and lower abundance relative to *Gluconobacter*.

**Figure 5.**
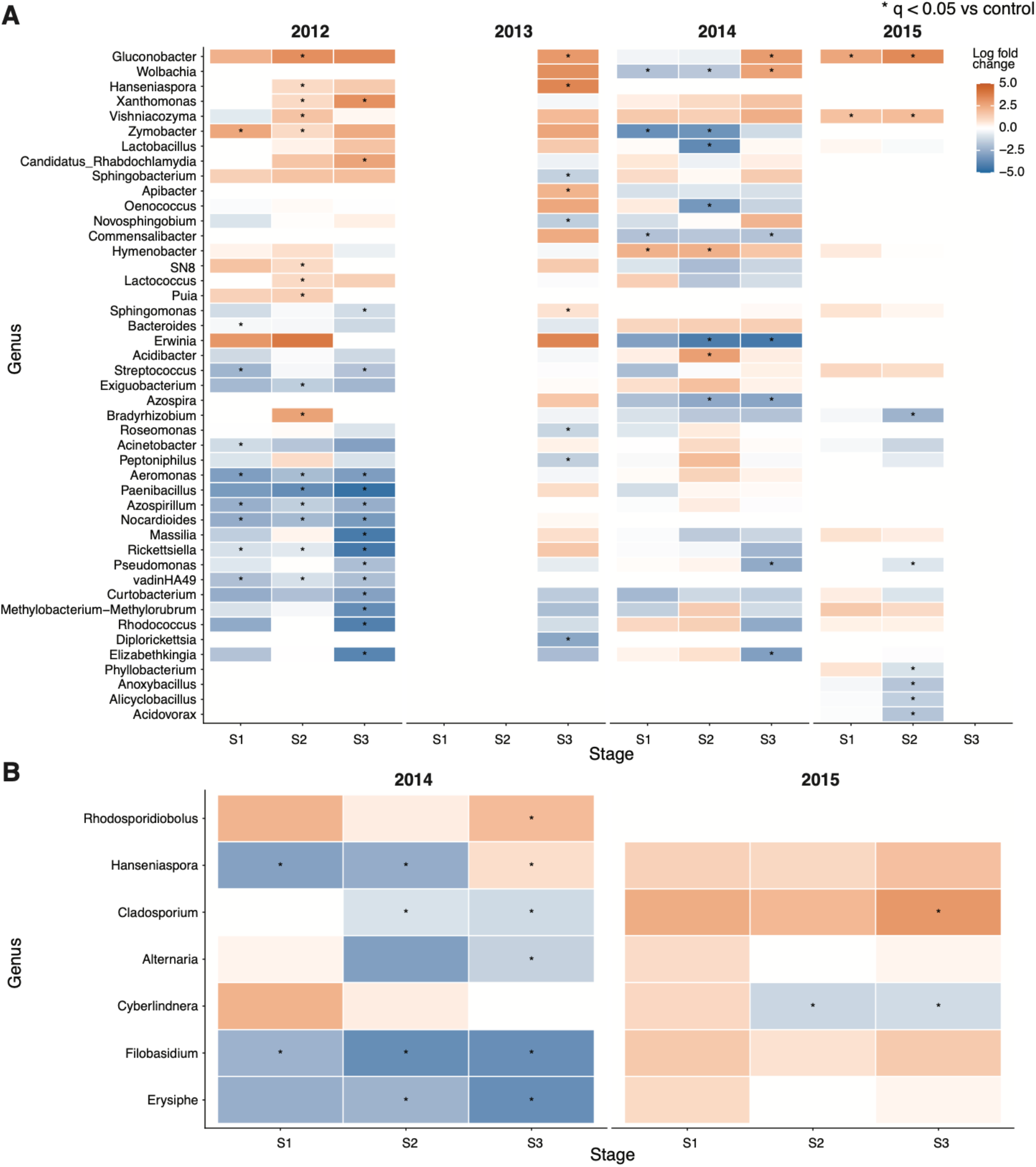
Differential abundance of bacterial (A) and fungal (B) genera across noble rot stages by year. Heatmaps show log fold changes (LFC) of genus-level relative abundance relative to control samples, calculated using ANCOM-BC. Positive LFC (orange) indicates enrichment relative to control; negative LFC (blue) indicates depletion. Asterisks indicate statistically significant differences (q < 0.05 after FDR correction). Analyses were performed separately for each vintage year. Only genera significant in at least one stage and year are shown. Fungal differential abundance analysis was not possible for 2012 and 2013 due to absence of fungal reads in control samples and insufficient control sample size, respectively.

The bacterial core microbiome (taxa present in ≥50% of samples within each stage) consisted of a single genus, *Pseudomonas*, across control, S1, and S2 stages (**Supplementary Fig. 2A**). At S3, the core microbiome expanded to include *Gluconobacter, Sphingomonas*, and *Burkholderia-Caballeronia-Paraburkholderia*, with *Elizabethkingia* appearing exclusively at S2. The emergence of *Gluconobacter* in the S3 core microbiome is consistent with its significant enrichment detected by ANCOM-BC across 2012-2014 vintages (**Fig. 5A**), reinforcing its role as a consistent and abundant member of the late-stage noble rot microbiome.

For fungal communities, differential abundance analysis was restricted to 2014 and 2015, as control samples in 2012 yielded near-zero fungal reads precluding the use of control as a reference group, and 2013 contained only a single control sample which did not meet the minimum sample size requirements for ANCOM-BC. In total, 8 unique fungal genera were identified across 22 significant genus-stage-year combinations (q < 0.05; Fig. 5B; **Supplementary Table S9**). In 2014, *Hanseniaspora* was significantly enriched at S1 and S3, while *Cladosporium* and *Alternaria* were significantly depleted at S3. *Filobasidium* was significantly depleted across all stages in 2014, and *Rhodosporidiobolus* was significantly enriched at S3. In 2015, *Cladosporium* was significantly enriched at S3, while *Cyberlindnera* was significantly depleted at S2 and S3. No fungal taxa formed a core community in control samples, consistent with low fungal biomass prior to *Botrytis cinerea* infection. *Hanseniaspora* was the only fungal genus present in the core microbiome across S1, S2, and S3, while *Lachancea* appeared exclusively at S3 (**Supplementary Fig. 2B**), consistent with the significant enrichment of *Hanseniaspora* at S3 observed in 2014. The overall pattern suggests a shift from filamentous fungi (*Cladosporium, Alternaria*) toward yeasts (*Hanseniaspora, Rhodosporidiobolus, Lachancea*) as noble rot progresses, consistent with the increasing sugar content and altered physicochemical conditions associated with advanced *Botrytis cinerea* infection.

### Acetic acid bacteria accumulate at advanced stages of noble rot across multiple vintages

Acetic acid bacteria (AAB) were detected across all noble rot stages, with total AAB abundance increasing dramatically at S3 in all three years with complete stage sampling (**Fig. 6A**). In 2012, total AAB relative abundance increased from less than 1% in control, S1, and S2 to a mean of ~10% at S3. In 2013, where only control and S3 were sampled, S3 reached a mean of ~25% total AAB abundance. In 2014, the highest AAB abundance was recorded, with S3 samples reaching a mean of ~37%. *Gluconobacter* was the dominant AAB genus across all years and stages, accounting for the majority of total AAB abundance at S3 (**Fig. 6B**). *Komagataibacter* was the second most abundant AAB at S3 in 2014, reaching ~7% mean relative abundance. *Kozakia* was consistently present at low levels across all stages and years. *Acetobacter, Gluconacetobacter*, and *Asaia* were minor components of the AAB community. The consistent and dramatic enrichment of AAB, particularly *Gluconobacter*, at S3 across three independent vintages strongly supports their role as indicators and potential agents of acetic acid production during advanced noble rot.

**Figure 6.**
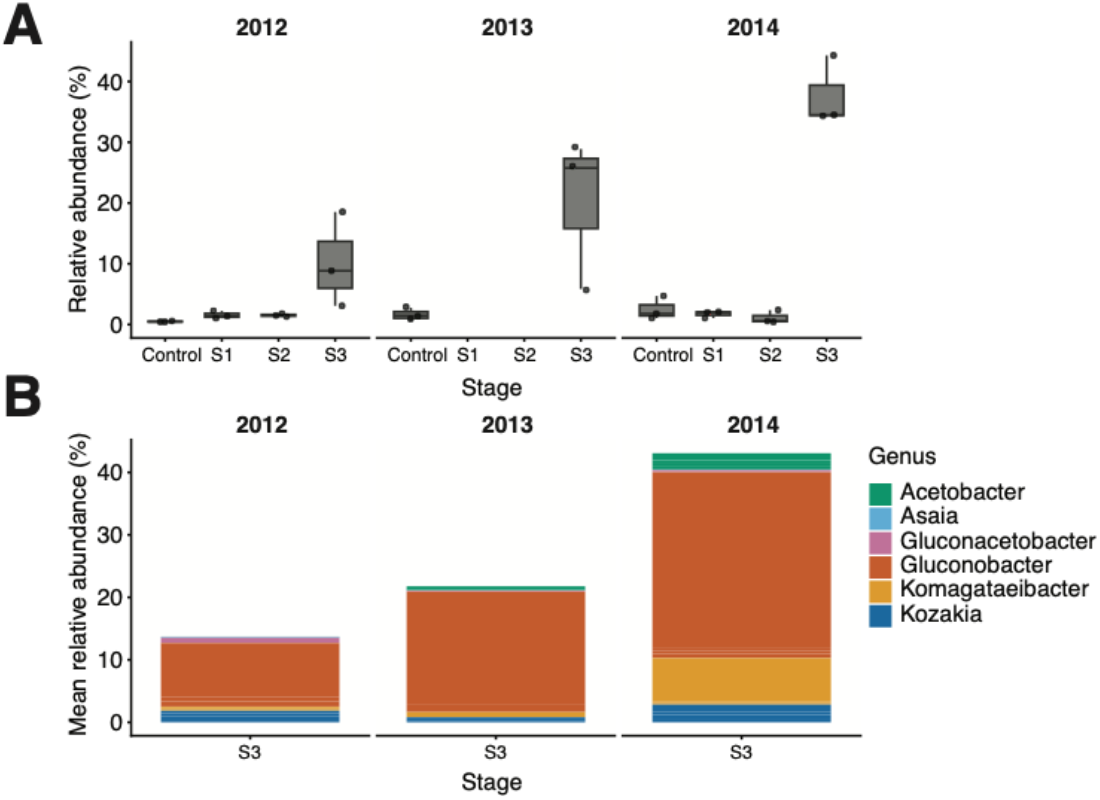
Acetic acid bacteria abundance and composition across noble rot stages. (A) Boxplots showing total relative abundance of acetic acid bacteria (Acetobacteraceae) per sample across noble rot stages in 2012, 2013, and 2014. Each point represents one sample. Colors indicate the noble rot stage. (B) Stacked barplots showing mean relative abundance of individual AAB genera at S3, the stage of peak AAB abundance, across the three vintages with complete stage sampling. Note that 2013 includes only control and S3 samples. 2015 is not shown due to the absence of S3 samples.

Co-occurrence network analysis based on Spearman correlations across 58 shared samples revealed that *Gluconobacter* is negatively correlated with *Pseudomonas* (**Fig. 7A**), consistent with their contrasting stage-specific abundances: *Pseudomonas* dominates early stages while *Gluconobacter* emerges at S3. *Kozakia* and *Komagataibacter* co-occurred positively with *Gluconobacter*, forming a coherent AAB cluster in the bacterial network. Cross-kingdom co-occurrence analysis revealed that *Gluconobacter* and *Komagataibacter* positively co-occurred with *Hanseniaspora* and *Lachancea* (**Fig. 7B**), two sugar-fermenting yeasts that also increase at advanced noble rot stages. *Elizabethkingia* showed a negative cross-kingdom correlation with *Lachancea*, consistent with its early colonizer behavior and decline at advanced infection stages.

**Figure 7.**
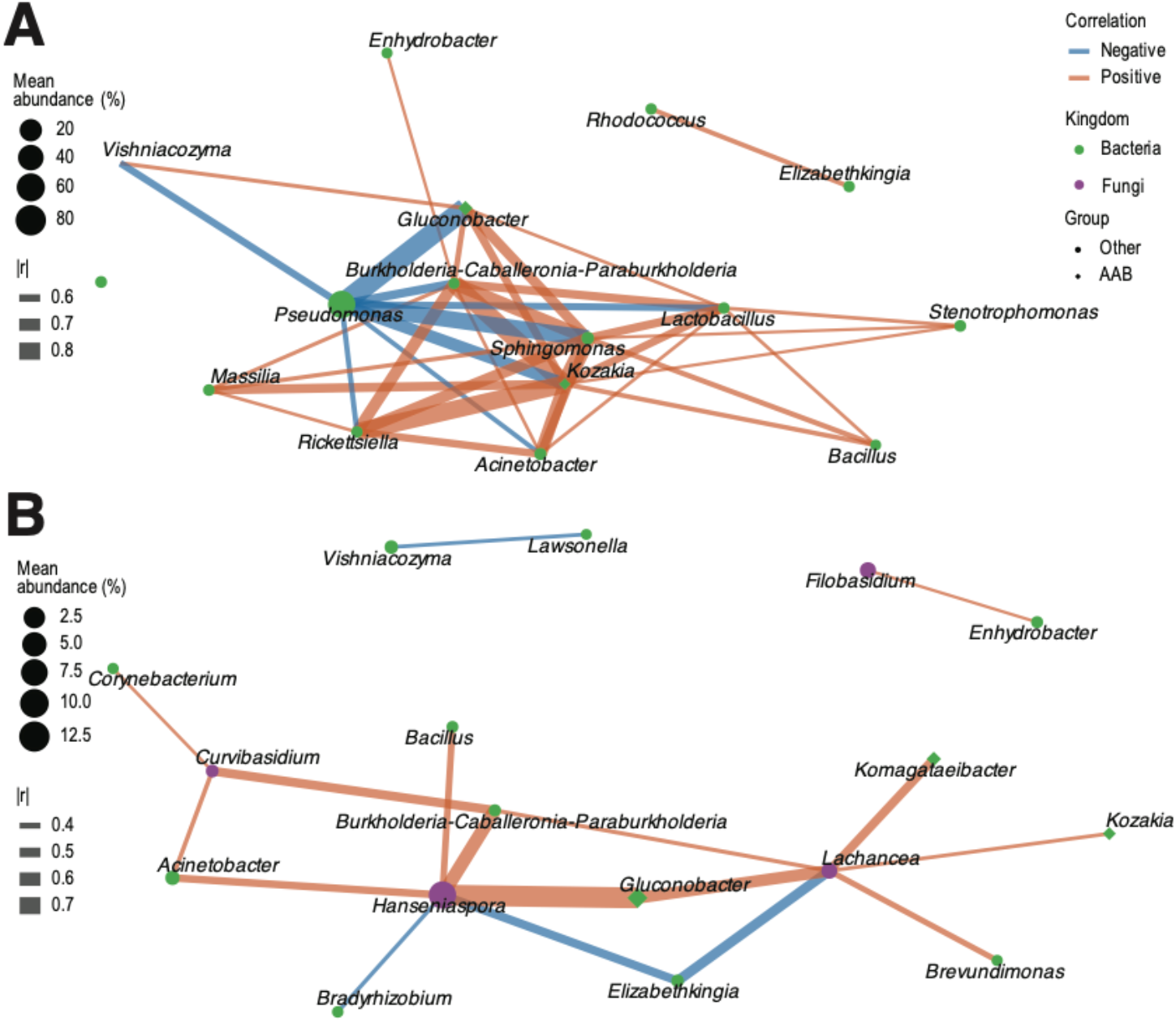
Co-occurrence networks of bacterial and cross-kingdom microbial associations on noble-rotted grape berries. (A) Bacterial co-occurrence network based on Spearman correlations across 58 samples shared between 16S and ITS datasets. (B) Cross-kingdom co-occurrence network showing significant correlations between bacterial and fungal genera. Nodes represent genera, colored by kingdom (green: bacteria, purple: fungi) and shaped by group (diamond: acetic acid bacteria, circle: other). Node size is proportional to mean relative abundance across all samples. Edge color indicates correlation direction (orange: positive, blue: negative) and edge width indicates correlation strength (|r|). Only significant correlations (q < 0.05, |r| ≥ 0.3 for cross-kingdom; |r| ≥ 0.5 for bacteria-only) after Benjamini-Hochberg correction are shown.

## Discussion

Noble rot is a distinct outcome of *B. cinerea* infection in which specific microclimatic conditions favor berry dehydration and compositional changes that support botrytized wine production. Despite extensive characterization of the physiological and metabolic changes associated with noble rot (Blanco-Ulate et al. 2015; Magyar 2011), the microbial communities colonizing berries across progressive infection stages have not been systematically characterized. Here, we show that noble rot drives a reproducible, stage-structured microbial succession across four consecutive vintages, marked by a shift from *Pseudomonas*-dominated bacterial communities toward enrichment of acetic acid bacteria at advanced infection stages, and a parallel transition in the non-Sclerotiniaceae fungal community from filamentous fungi toward fermentative yeasts.

*Pseudomonas* spp. dominated the bacterial communities consistently across control, S1, and S2 stages in all four vintages. *Pseudomonas* spp. are among the most common bacterial colonizers of plant surfaces, including grape berries, where they have been reported as prevalent phyllosphere inhabitants across multiple varieties and geographic origins (Barata et al. 2012; Bokulich et al. 2014; Wei et al. 2018; Zarraonaindia et al. 2015). Their prevalence on healthy and early infected berries likely reflects their generalist lifestyle and ability to exploit the nutritional resources available on intact or mildly damaged berry surfaces. The decline in *Pseudomonas* relative abundance at advanced stages is consistent with competitive displacement by the expanding AAB community rather than an absolute reduction in *Pseudomonas* abundance, though absolute quantification would be needed to confirm this. The negative co-occurrence between *Pseudomonas* and *Gluconobacter* in the network analysis supports this interpretation. Whether this reflects direct antagonism, niche displacement driven by changing substrate chemistry, or both remains to be determined.

AAB, particularly *Gluconobacter*, were reproducibly enriched at advanced noble rot stages. *Gluconobacter* was significantly enriched at S3 in all vintages with complete stage sampling (2012, 2013, and 2014; q < 0.05) and was the only AAB genus to join the bacterial core microbiome exclusively at this stage. Total AAB abundance reached means of approximately 10%, 25%, and 37% of the bacterial community at S3 in 2012, 2013, and 2014, respectively. These findings are consistent with culture-based studies reporting substantial AAB population increases on *B. cinerea*-damaged compared to healthy berries (Barata et al. 2012), and extend them by demonstrating that this enrichment is stage-specific, reproducible across vintages, and dominated by *Gluconobacter* rather than *Acetobacter*, which is more commonly reported as the primary AAB in grape sour rot (Hall et al. 2018). Whether this difference reflects the distinct substrate chemistry of noble rot versus sour rot berries, particularly the higher sugar and lower ethanol content at harvest, or other ecological factors, remains to be determined.

*Gluconobacter* is distinguished from *Acetobacter* by its greater efficiency in oxidizing sugars to ketoacids and gluconic acid, while *Acetobacter* more efficiently oxidizes ethanol to acetic acid (Raspor and Goranovic 2008). The high sugar content of S3 noble rot berries may therefore selectively favor *Gluconobacter*. Gluconic acid is a commonly used as a marker of *B. cinerea* infection severity in must assessment (Barata et al. 2012; Magyar and Soós 2016), and while *B. cinerea* itself produces gluconic acid via glucose oxidase activity; *Gluconobacter* likely contributes to its accumulation in heavily infected berries. *Komagataibacter*, the second most abundant AAB at S3 in 2014 (~7% of the bacterial community), includes cellulose-producing species that can form biofilms on berry surfaces and complicate pressing during winemaking (Raspor and Goranovic 2008), though whether cellulose production occurs under noble rot conditions requires direct confirmation.

Sclerotiniaceae dominated the fungal community across all stages and vintages, confirming widespread *B. cinerea* colonization even at early infection stages. The inability to resolve Sclerotiniaceae to genus level using the UNITE database likely reflects the limited discriminatory power of the ITS region for closely related taxa within this family, including *Botrytis* spp., which cannot be reliably distinguished at the species level using ITS alone (Holst-Jensen et al. 1997; Staats et al. 2005). Beyond Sclerotiniaceae, the fungal community excluding this family showed a transition from filamentous fungi, particularly *Cladosporium* and *Alternaria*, toward fermentative yeasts including *Hanseniaspora* and *Lachancea* at advanced infection stages. Cladosporium and Alternaria are common fungal colonizers of grape berry surfaces (Barata et al. 2012), and their depletion at S3, confirmed by ANCOM-BC in 2014, may reflect environmental filtering and a shift favoring fermentative yeasts under the high-sugar, low water activity conditions of heavily colonized berries, though the mechanisms remain speculative.

*Hanseniaspora* was among the most consistently abundant yeasts across noble rot stages, present in the core microbiome from S1 onward and significantly enriched at S3 in 2014. *Hanseniaspora uvarum* is among the dominant yeasts on ripe grape berries globally and an important early fermentation contributor (Barata et al. 2012; Jolly et al. 2014). Its consistent presence across infection stages suggests it colonizes berries independently of infection status but proliferates under the conditions of advanced noble rot. *Lachancea thermotolerans*, which appeared in the core microbiome exclusively at S3, is recognized for its capacity to produce lactic acid and modulate wine acidity (Benito 2018), and its emergence at advanced stages may reflect adaptation to the high-sugar and chemically altered environment of S3 berries.

The positive co-occurrence between *Gluconobacter* and the fermentative yeasts *Hanseniaspora* and *Lachancea* suggests concurrent enrichment of AAB and yeasts at advanced noble rot stages. Both groups are favored by the high sugar conditions associated with damaged berry tissue and commonly proliferate in heavily colonized grape berries (Barata et al. 2012). Their co-occurrence likely reflects shared ecological preferences, though metabolic interactions such as yeast-produced ethanol serving as substrate for AAB-mediated acetic acid production cannot be excluded and may contribute to the elevated volatile acidity often associated with botrytized must (Magyar and Soós 2016). The pre-harvest assembly of this community has practical implications: the composition and relative abundance of AAB and fermentative yeasts at harvest are likely to influence the microbial inoculum entering fermentation, must chemistry, and the *Saccharomyces* establishment. Bacterial diversity in botrytized wine fermentations from the same vineyard was substantially higher than in fermentations of healthy grapes, and AAB were strongly suppressed following *Saccharomyces* inoculation, highlighting how the field-assembled microbial communities can shape fermentation microbiology (Bokulich et al. 2012).

Sampling across four consecutive vintages allowed us to distinguish reproducible, stage-associated patterns from vintage-specific variation. The enrichment of *Gluconobacter* at S3, the dominance of *Pseudomonas* at early stages, and the shift from filamentous fungi to yeasts were consistent across years, suggesting deterministic responses to the physiological changes associated with *B. cinerea* infection. Vintage effects were nonetheless detectable: year was a significant factor in bacterial community composition by PERMANOVA, and the magnitude of AAB enrichment at S3 varied substantially across vintages. These differences likely reflect vintage-specific variation in infection progression, weather conditions, and berry chemistry, but their specific drivers remain to be identified.

Interpretation of these results should nonetheless consider several methodological limitations. Because amplicon sequencing provides relative abundance rather than absolute quantification, the observed enrichment of AAB at advanced noble rot stages cannot be directly translated into population sizes, nor can active and inactive community members be distinguished. Culture-based assays or targeted qPCR approaches would therefore be needed to confirm absolute AAB population dynamics. In addition, fungal differential abundance analysis could only be performed for the 2014 and 2015 vintages, owing to the near absence of non-*Botrytis* fungal reads in 2012 controls and insufficient control sample size in 2013. Low sequencing depth also led to the exclusion of 2015 S3 bacterial samples, limiting characterization of late-stage bacterial dynamics in that vintage. Finally, the co-occurrence network represents correlations aggregated across all samples and stages and therefore does not resolve stage-specific microbial interaction dynamics.

This study provides a comprehensive characterization of the grape berry microbiome across progressive stages of noble rot. *B. cinerea* infection is associated with a reproducible microbial succession in which AAB, particularly *Gluconobacter*, and fermentative yeasts are enriched at advanced stages, whereas early colonizers, such as Pseudomonas and filamentous fungi, decline. The consistency of this succession across four vintages points to a reproducible ecological process associated with the progressive physiological transformation of the noble-rotted berries. These findings have direct implications for botrytized wine production, as the pre-harvest microbiome influences must chemistry, fermentation, and wine quality. Future work should address the metabolic interactions between AAB and yeasts on noble-rotted berries, the drivers of vintage variation in AAB abundance, and whether microbiome composition at harvest predicts fermentation outcomes.

## Data availability

The sequencing data generated for this project are available at NCBI (BioProject: PRJNA1468206; https://www.ncbi.nlm.nih.gov/bioproject/PRJNA1468206). Code is available at https://github.com/CantuLab/Noble-rot-microbiome

