## Supplementary Figures for "*Botrytis cinerea* infection reshapes the grape berry microbiome during noble rot"

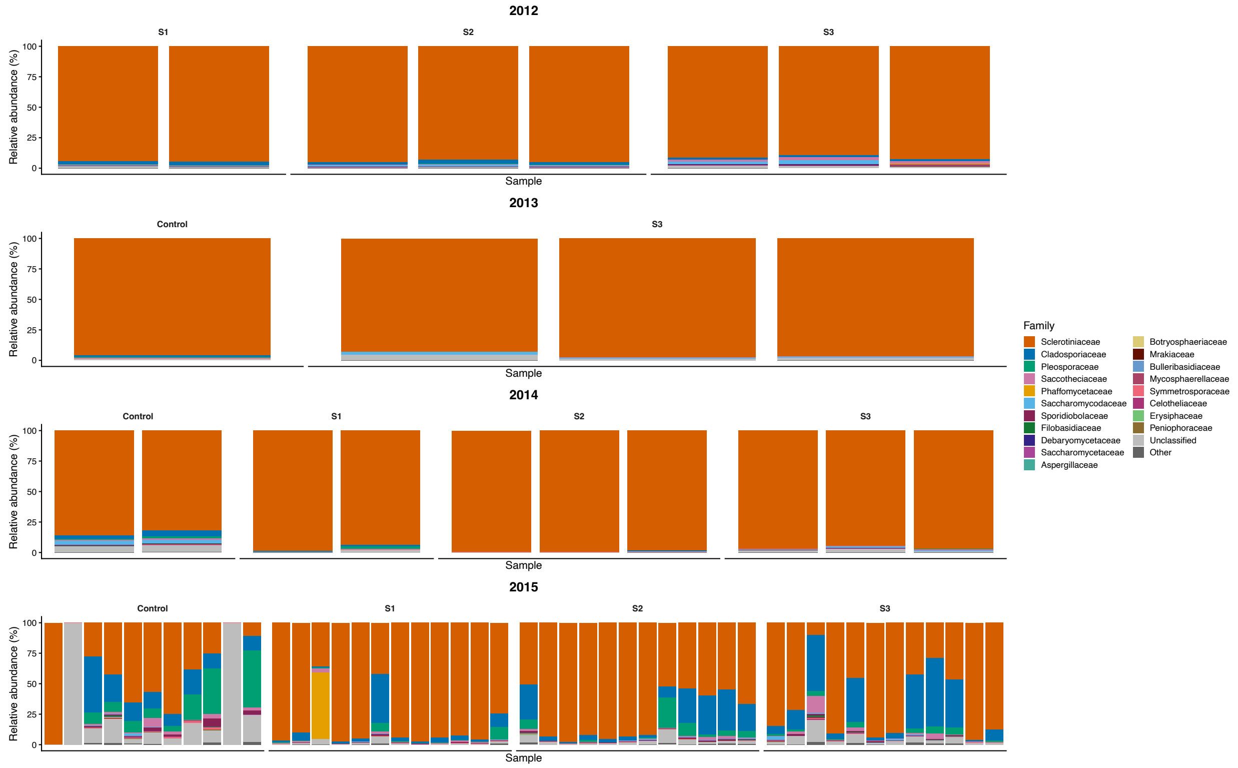


**Supplementary Figure 1. Fungal community composition at family level across noble rot stages and vintages.** Stacked barplots show the relative abundance (%) of the top 20 fungal families (ITS1 region) in each sample, grouped by noble rot stage (Control, S1, S2, S3) and vintage year (2012, 2013, 2014, 2015). Each bar represents one sample. Families not in the top 20 are grouped as "Other". Sclerotiniaceae, representing primarily unclassified *Botrytis cinerea*, dominates the fungal community across all stages and vintages. Note that 2012 lacks control samples due to near-zero fungal reads in control berries, and 2013 includes only control and S3 stages. Fungal taxonomy was assigned using the UNITE v10.0 database.

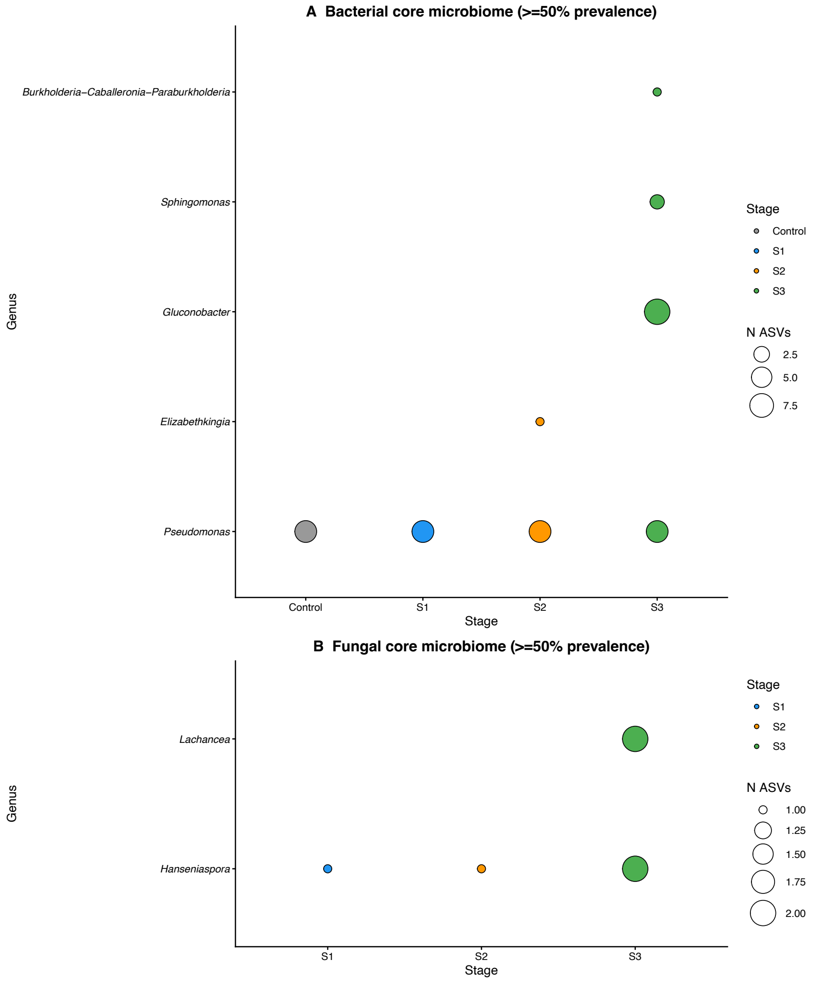


**Supplementary Figure 2. Core microbiome of bacterial (A) and fungal (B) communities across noble rot stages.** Bubble plots show genera present in at least 50% of samples within each noble rot stage (Control, S1, S2, S3), based on 16S rRNA gene (bacteria) and ITS1 (fungi) amplicon sequencing across all vintages combined. Bubble size represents the number of ASVs assigned to each genus within the core.
